# How cells align to structured collagen fibrils: A hybrid cellular Potts and molecular dynamics model with dynamic mechanosensitive focal adhesions

**DOI:** 10.1101/2024.07.10.602851

**Authors:** Koen A.E. Keijzer, Erika Tsingos, Roeland M.H. Merks

## Abstract

Many mammalian cells, including endothelial cells and fibroblasts, tend to align and elongate with the orientation of extracellular matrix (ECM) fibers in a gel when cultured in vitro. During cell elongation, clusters of focal adhesions (FAs) form near the poles of the elongating cells. FAs are mechanosensitive clusters of adhesions that grow under mechanical tension due to the cells’ pulling on the ECM, and shrink when the tension is released. Using a mathematical modeling approach, we study the hypothesis that reciprocity between cells and the ECM drives cell shape changes. We show that FAs are preferentially stabilized along the orientation of ECM fibers, where the cells can generate more tension than perpendicular to the ECM fibers. We present a hybrid cellular Potts model that represents the ECM as an off-the-lattice network of cross-linked deformable fibers, whereas the cell is represented on the lattice. Multiple FAs are modeled individually by an independent rate of FA assembly and a mechanoresponsive FA disassembly. The resulting computational model predicts stiffness-dependent cell spreading and local ECM remodelling, and ECM-alignment dependent cell elongation. The effects combined suffice to explain how cell morphology is determined by local ECM structure and mechanics.

## 1 Introduction

The extracellular matrix (ECM) plays a crucial role in development and in disease. For example, the ECM plays a role in cancer cell migration (Najafi et al., 2019; Yamaguchi et al., 2005), wound healing (Maquart and Monboisse, 2014; Diller and Tabor, 2022) and angiogenesis (Stupack and Cheresh, 2002). The ECM is a complex collection of large fibers such as collagen, fibronectin and other proteins (Theocharis et al., 2016). The orientation of fibers in the ECM is thought to play an important role in tumor vascularisation (Balcioglu et al., 2016), mechanical cell-cell communication (Nahum et al., 2023), and blood clot formation (Kim et al., 2017b). As a dynamic and complex environment, the ECM is continuously remodeled by cells both chemically, through the synthesis and degradation of fibers, and mechanically, by pulling and reorienting fibers (Theocharis et al., 2016; Winkler et al., 2020). Consequently, as cells react to fiber orientation and reorient fibers, studying this interplay is crucial to fully understand the mechanical reciprocity between cells and the ECM. In this paper, we introduce a computational model that demonstrates how mechanical reciprocity emerges from simple assumptions on the adhesion between cell and matrix.

In vitro studies examining cell responses to fiber orientation and other mechanical ECM properties often involve the use of artificial cell cultures on fabricated substrates. The morphology of cells varies widely depending on substrate stiffness and geometry. Soft substrates result in small cells, whereas cells increase their area on stiffer substrates. Many cell types, such as endothelial cells, fibroblasts, smooth muscle cells and osteogenic cells, show a monotonic increase in spreading with substrate stiffness, achieving maximum spreading on very stiff substrates such as coated glass (Yang et al., 2017). Other cell types, such as T-cells, show a biphasic spreading response to substrate stiffness, showing maximal cell spreading at an intermediate optimal level of substrate stiffness (Janmey et al., 2020; Wahl et al., 2019). Next to studies on the effect of substrate stiffness on cell area, there have been many studies elucidating the effect of substrate geometry on cell morphology, for example using pillared substrates (Feld et al., 2020) or micropatterned substrates (D’Arcangelo and McGuigan, 2015). Here we focus on those studies fabricating anisotropic fibrous substrates, showing that many cell types tend to align with the fibers in the substrate on which they are cultured (Lai et al., 2012; Friedrichs et al., 2007; Sapudom et al., 2023; Chaubaroux et al., 2015). Both the mechanical properties of the fibers and the cell adhesive properties influence cell elongation. Cells, for example, do not elongate on fragile fibers as the fibers rupture due to cellular forces (Friedrichs et al., 2007). Additionally, if the cell lacks the proper ECM receptors, they do adhere but cannot elongate along the fibers (Friedrichs et al., 2007). In short, cells sense both ECM stiffness and the orientation of fibers in the ECM.

Cells do not only respond to cues in the ECM, such as collagen fiber orientation and overall ECM stiffness, they also restructure the ECM by exerting force on it. On aligned collagen substrates, cells recruit and bundle collagen fibrils on their leading edge (Friedrichs et al., 2007). Contractile cells deform fibrous matrices, and reorient the fibers to point towards the cell (Kim et al., 2017a). Hence, cells can restructure their micro-environment by mechanical forces.

Clearly, mechanical reciprocity, the mechanical two-way interaction between the cell and the ECM, is a complex subject combining cell biology and mechanics, requiring an understanding of ECM mechanics cell behavior and of cell behavior. In general, computational models can give insight into complex mechanisms and there is a wide variety of work on computational models of cell-ECM interaction (Crossley et al., 2024). Here we briefly review a selection of specific computational models that capture the fibrous nature of the ECM and model the cell-ECM interaction through mechanosensitive adhesions; furthermore we highlight if mechanoreciprocity emerges, and if so the two factors that are crucial to be included in a computational model for studying the reciprocal effect between fiber alignment and cell morphology namely (i) ECM mechanics, and (ii) dynamically changing cell morphology. We view these two factors as the two aspects of mechanical reciprocity.

Computational models of extracellular matrix (ECM) mechanics frequently simplify cell morphology to basic geometric shapes, yet they enable the emergence of complex behaviors by integrating mechanosensing mechanisms through cellular adhesions. In one computational model of a realistic 3D finite element (FE) representation of the ECM, it was shown how durotaxis, an example of cell mechanosensing where the cell migrates upwards stiffness gradients, emerges from cell contractility and force dependent adhesion to the ECM (Paukner et al., 2023). In this model, the cell deforms the ECM, but no changes to the cell’s morphology are possible as the cell is modeled as a point particle with an adhesive annulus. In another model of cell migration, durotaxis likewise emerges from cellular contractility, and mechanosensitive adhesion without using a FE simulation for the ECM but rather a simplified bead-spring network that deform under the contractile forces of a rotating ellipodial cell (Feng et al., 2019). It is noteworthy that different implementations of similar conceptual models of cell-ECM interaction support the notion that the interplay of cytoskeletal force and mechanosensitive adhesion are important factors in durotaxis (Paukner et al., 2023; Feng et al., 2019; Rens and Merks, 2020; Mathieu et al., 2024). Next to the mechanosensing of substrate stiffness, cells can sense the fiber orientation and migrate effectively along fibers (Feng et al., 2019) in a process called contact guidance (SenGupta et al., 2021). Interestingly, in silico, the cells ability to sense fiber orientation disappears if the fibers bending modulus is too high (Feng et al., 2019), suggesting that the cell probes this information through mechanically interacting with the fibers, again suggesting that mechanosensitive adhesions and cell contractility are important factors. Other studies focused more on ECM remodeling by the cell than on cell migration. In non-fibrous ECMs, the matrix typically strains under cellular contractile forces, whereas on fibrous ECMs the cell restructure and reorient the matrix, leading to accumulation and reorientation of fibers. In silico models suggest that cell-mediated fiber accumulation can enhance cell spreading by increasing the number of available bindings sites for the cellular adhesion (Cao et al., 2017). The computational studies that we discussed thus far, although giving insight in mechanical cell-ECM mechanism, did not take the dynamic cell’s morphology into account. Including the morphology of the cell in a computational model opens the doors to understanding more involved mechanical mechanisms as, for example, different cell migration modes can emerge based on adhesion maturation and stress fiber strength in a 3D finite element model of a moving cell on a non-fibrous uniform ECM (Vargas et al., 2020). One of the first models that uses both discrete fibrous ECM and a deformable cell that was used to study cell-ECM interactions was used to explain how bands form between two contractile cells in a fibrous ECM, and how the two cells elongate towards each other by the remodeled matrix (Reinhardt et al., 2013; Crossley et al., 2024).

To better understand the effect of ECM orientation on cell morphology, both deformable fibers ECM and dynamic cell morphology should be included in a computation model. A model that is effective in modeling complex cell shapes is the cellular Potts model (CPM) (Graner and Glazier, 1992). Although the CPM has primarily been applied in multicellular development (Hirashima et al., 2017), more recently it has been employed for understanding single cell mechanics (Albert and Schwarz, 2016; Scianna and Preziosi, 2021; Schakenraad et al., 2022). The CPM has been coupled with mechanical ECM models, forming a hybrid CPM, to study the effect of the ECM on cell shape, for instance explaining the role of cell contractility on cell elongation when the ECM is stretched (Rens and Merks, 2017), or investigating the role of mechanical cell-cell communication during vasculogenesis (van Oers et al., 2014). These previous models use similar mechanical models for the ECM, namely homogeneous elastic materials, that do not capture plastic deformations or anisotropic strain response to the cell contractile forces observed in fibrous networks. Incorporating plastic deformations and anisotropic strain responses into a hybrid CPM can be achieved using coarse-grained molecular dynamics simulations that approximate the ECM, employing beads linked together with springs to form a collagen-like network (Tsingos et al., 2023). In short, the CPM is a flexible model that can captures different cell morphologies and allows coupling with external ECM models.

The choice on how to couple cellular morphology and ECM dynamics is delicate, as this choice encodes the biological hypothesis of how cells sense and react with ECM. Early CPM-ECM couplings assumed durotaxis to be the driving force where cellular protrusions are stabilized on highly stressed substrates (van Oers et al., 2014). Subsequently, this coupling was extended by adding a comprehensive model of the mechanosensitive adhesions between the cell and the ECM, leading to emergent cell spreading, spontaneous cell elongation, and durotaxis (Rens and Merks, 2020). We applied the coupling between cell and ECM proposed by Rens and Merks (2020) and apply it to a hybrid CPM with discrete fibrous ECM (Tsingos et al., 2023). The coupling is made by assuming that the cell exerts cytoskelletal contraction forces through integrin based adhesions that behave according to the two-spring model (Schwarz et al., 2006; Doyle et al., 2022). In essence, the two-spring model views an adhesion as mediator between the contractile forces of the cell against the restoring forces of the ECM, and hence sees the tension on the adhesion build-up slowly on soft ECMs and quickly on stiff ECMs as the cell applies its contractile forces. Additionally, it is assumed that adhesions strengthen as tension increases, which is quantified by the number of integrin proteins bound to the adhesion. These assumptions, when combined with an isotropic material in a hybrid CPM, are sufficient to produce phenomena such as cell spreading, spontaneous elongation, and durotaxis (Rens and Merks, 2020). In this paper, we implement this two-spring adhesion model in the hybrid CPM with discrete fibrous ECM (Tsingos et al., 2023), and we use the new model to investigate the interplay between fiber orientation and cell morphology. Specifically, with this new fibrous ECM model, we show how cell elongation on oriented gels can be considered a special case of stiffness dependent cell spreading, as fibers are easier to bend than to stretch. Furthermore, we show how cell protrusions reorient fibers, thereby increasing tension on adhesions and stabilizing the protrusions.

## 2 Methods

### 2.1 Modeling Approach

We have introduced dynamic descriptions of mechanosensitive focal adhesions (FA) into a previous, hybrid cellular Potts (CPM) and molecular dynamics (MD) model (Tsingos et al., 2023). The CPM part dynamically describes cell shape changes, and the MD-part simulates a cross-linked network of ECM fibers, and its dynamical response to cellular forces. In our previous work the CPM was connected to the MD through static adhesion particles. In the present model the build-up and break-down of FA’s are modeled dynamically, using an ordinary differential equation model that describes FAs as clusters of integrins whose breakdown rate is assumed to be dependent on the mechanical tension within the FAs Novikova and Storm (2013), as in one of our previous models featuring a continuum description of the ECM (Rens and Merks, 2020). Figure 1(A) provides a diagram of the key elements of the hybrid model and the interactions between its components.

**Figure 1:**
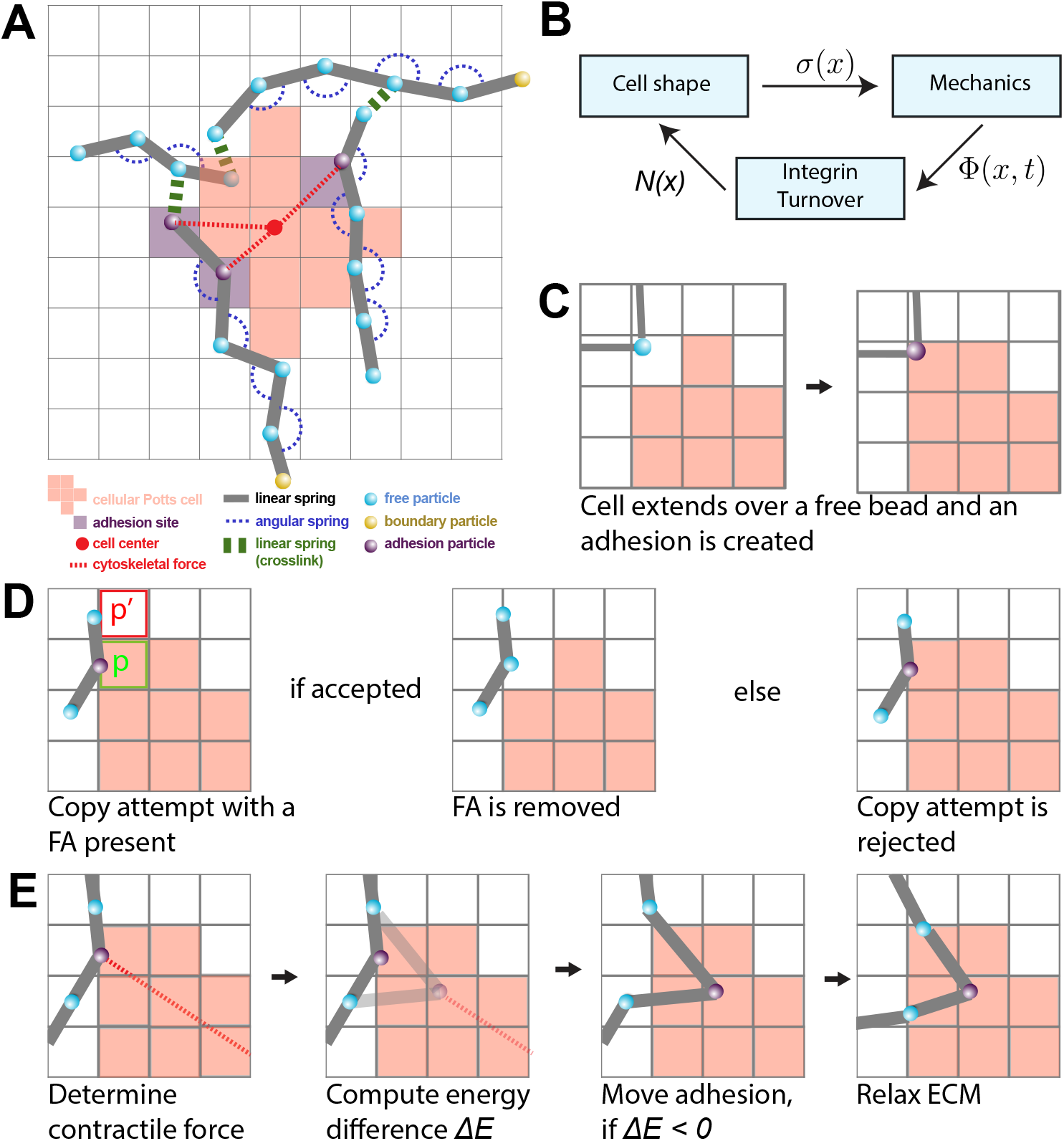
A: A schematic overview of the components in the model. The lattice bound cellular Potts cell (red squares) is attached to the beads (blue) and springs (grey) of the ECM at FAs (purple). The contractile force (dashed red) is visualized between the FAs and the center of mass of the cell (red point). The FAs connect the cell to the strands of the ECM. The strands are cross-linked (green) and particles outside of the simulation domain are fixed (yellow). B: The different submodels of the model are linked together as shown in this figure. The CPM is responsible for the cell shape, and sends the lattice *σ*(*x*) to the mechanical part of the model. The mechanical part computes the tensions Φ(*x*) in the FAs and gives this to the FA part of the model. The tensions are used to calculate the number of bound integrins *N*(*x*) in each FA. These numbers are then used in the CPM to update the new cell shape. C: A scheme showing the creation of a new FA. When a cell extends over a free bead, it becomes an adhesion bead. D: Scheme showing a retraction copy-attempt when a FA is present. The FA is removed if the copy1-a1ttempt is accepted (left). The presence of the FA increases the likelihood of rejecting the copy-attempt (right). E: a FA is pulled towards the center of the cell in a process resembling a CPM copy-attempt: First the energy change between the original position of the FA and the new FA is determined by only considering springs that are directly attached to the FA. The FA is moved if the energy difference is negative. Finally, the complete ECM is relaxed by running the molecular dynamics simulation.

The models are coupled using an operator splitting approach. The three submodels sequentially iterated to steady state, such that the output state of one submodel is used as the input state for the next submodel (Figure 1(B)). The simulations were run until quasi steady state where we did not observe large further changes. In the remainder of this Section, we will describe each of the three submodels as well as the coupling strategies.

### 2.2 Cellular model

To describe cell shape changes, we make use of the cellular Potts model(CPM) (Graner and Glazier, 1992; Hirashima et al., 2017). The CPM is a lattice based model where cell shape is defined as a collection of connected lattice sites. We implemented a CPM on a square grid Λ of 200 × 200 lattice sites. Each lattice site 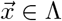 is assigned a spin σ(*x*) ∈ **Z**_≥0_ which defines a spin field *σ* : Λ → **Z**_≥0_. The collection of connected lattice sites that have the same positive spin *n* define the shape of cell *n*. The set of lattice sites with spin 0 are not occupied by a cell. The CPM evolves through a sequence of random extensions and retractions, whose probability is given by a balance of contractile, extensile and forces due to adhesion with the ECM. These are given by a Hamiltonian energy function,

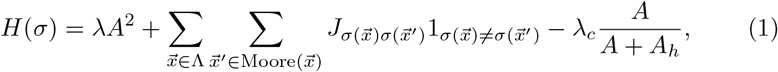

where *A* is the area of the cell, Moore 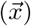 is the set of lattice sites in the Moore neighborhood of 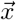, and *λ, J, λ*_*c*_, and *A*_*h*_ are parameters. The first part of Equation (1) describes the contractile force of the cell with rate *λ*. The second term penalizes, with strength *J*, interfaces between the cell and the medium, effectively creating a line tension along the cell’s perimeter. The final term describes the formation of non-integrin based FA with the substrate that bind with strength *λ*_*c*_ with a saturation parameter *A*_*h*_.

The Hamiltonian is minimized through Metropolis dynamics as previously described (Graner and Glazier, 1992), thus dynamically updating the cell’s shape. Briefly, we iteratively select a random lattice site 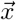 and a random, adjacent lattice site 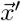. We then calculate the energy difference Δ*H* that would occur due to the update, and accept the copy attempt with probability *P*(Δ*H*) = 1 if Δ*H <* 0, and *P*(Δ*H* = exp(−Δ*H/T*) for Δ*H* ≥ 0, where *T* is a cell motility parameter.

The acceptance probability of a copy-attempt is determined by the energy change

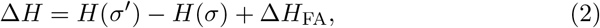

where Δ*H*_FA_ is an additional penalty for breaking integrin bonds that the cell might have with the ECM at that location.

The term Δ*H*_FA_ in Equation (2) is non-zero only if the copy attempt would be a retraction from a site 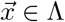 that contains a FA and in this case,

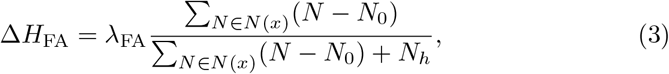

where 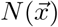 is the set with elements the number of integrins in a FA situated at 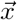, *λ*_*F A*_ a scaling parameter, *N*_0_ the initial size of a FA, and *N*_*h*_ a saturation parameter. If a copy attempt is accepted that leaves a FA outside of the cell, the FA is removed as in Figure 1(D). If a copy attempt is accepted that extends over a free bead of the ECM, then a new FA is created as in Figure 1(C).

### 2.3 Extracellular matrix model

The ECM is built out of beads with positions 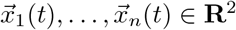. which are governed by the overdamped Langevin equation of motion

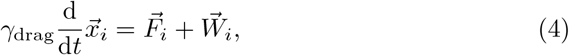

with *γ*drag a drag coefficient and *F*_*i*_ the force on the i-th particle and 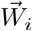 a random force satisfying 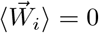 and 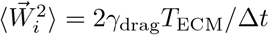 with *T*_ECM_ a parameter for degree of noise in the system and Δ*t* the size of a timestep. Equation (4) was integrated to a steady-state, with fixed Δ*t* during the simulation using the HOOMD-blue molecular dynamics libary (Anderson et al., 2020).

The forces on the beads are defined by harmonic potentials. If a bead pair (*i, j*) is linked with a spring of spring constant *k* and rest length *r*_0_ then the potential is given by

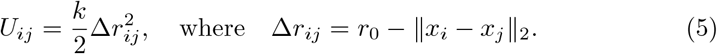

Likewise we add a bending rigidity to a triple of beads (*i, j, k*) by adding in a harmonic potential for the the angle

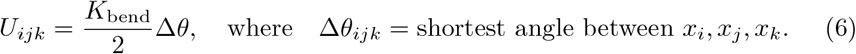

where *K*_bend_ is the bending rigidity. The exact linking of beads depends on the type of ECM that is studied. Some beads are fixed in space and are excluded from equation (4). These beads are at the boundary of the system, effectively clamping the ECM at the sides of the integration domain.

The ECM is described as a set of fibers connected through cross-linkers, thus forming an fiber network. A fiber is built out of *n*_beads_ beads which are linked together with springs of stiffness *K*_polymer_ and rest length *r*_polymer_. The contour length of strand is then *r*_polymer_ · (*n*_beads_ − 1). Consecutive triplets of beads in a fiber are connected with a harmonic potential with bending rigidity *K*_bend_, ensuring that unforced fibers remaing straight. To create a network, cross-linkers are added to the fibers. Cross-links are defined as springs with a small rest length and stiffness equal to *K*_cross_ = *K*_polymer_ and link different fibers together to from a connected network.

To create a fiber network, we followed the Method introduced in Tsingos et al. (2023) with small modifications for creating networks of aligned fibers. Briefly, we distributed *N*_strands_ randomly and uniformly in space, selecting fiber orientations from the Von Mises distribution to control the degree of fiber alignment. Fibers were created as follows: The position of the middle bead 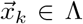 with *k* := floor(*n*_beads_*/*2) of a strand was selected at random from a uniform distribution, and a random angle *θ* ∈ [0, 2Π] was selected from the Von Mises distribution with *μ* ∈ [0, 2*π*) and *κ* ∈ [0, ∞). Then the remaining positions 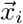 making up the beads where defined via

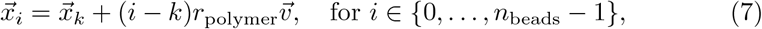

where *v* = (cos *θ*, sin *θ*) is a unit vector with angle *θ*. Constructing the fiber positions in this way ensures that the middle of each fiber is within the simulation domain, while only the endpoints might extend beyond the simulation domain. After fibers have been introduced, the network is cross-linked using the Method used in previous work (Tsingos et al., 2023).

### 2.4 Focal adhesions

FAs are modeled as clusters of catch slip bonds (Novikova and Storm, 2013; Rens and Merks, 2020; Schwarz et al., 2006). The cluster is in constant flux as integrins bonds are added and removed from the cluster. The integrin addition rate is independent, whereas the removal rate is suppressed by mechanical tension due to the contractile force of the cell and the restoring force from the ECM. The number of integrins *N* in a single focal adhesion under tension Φ is computed using the equation

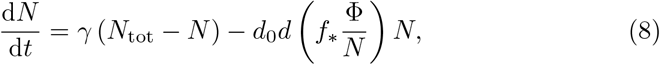

where *γ* is the binding rate of integrins, *N*_tot_ is the maximum number of integrins in a single FA, *d*_0_ a base detachment rate, *f*_∗_ a force scale, and *d*(*φ*) is a function of the tension per integrin that encodes the response of mechanical tension to the unbinding rate of a single integrin. We use a model for a catch-slip integrin that takes the form

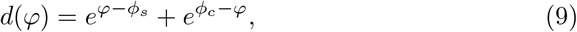

where *ϕ*_*s*_ and *ϕ*_*c*_ describe the slip and catch regime of an integrin (Novikova and Storm, 2013; Rens and Merks, 2020).

The tension Φ on the FAs is due to the force balance of the contractile, cellular force and the resultant force from the ECM. To calculate the contractile force we apply a model that force is proportional to the distance to the cell center (Lemmon and Romer, 2010), effectivily modeling the cytoskeleton as a spring connecting the FA with the cell’s center. The energy that the cytoskeleton exerts is then assumed to be 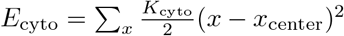 with *x* the position of a FA and *x*_center_ the center of the cell to which the FA belongs and with *K*_cyto_ a spring constant encoding the isotropic cell force. Likewise the energy of parts of the ECM that is directly linked to the FA is defined as *E*_ecm_. a FA is now moved as follows: First the total energy is computed *E* = *E*_cyto_ + *E*_ecm_. Then, the energy *E*^*′*^ is computed if the FA was to move one lattice site towards the center of the cell. If the difference *E*^*′*^ − *E* is negative, the FA is moved to the new position and otherwise is kept in place.

### 2.5 Parameter values

Parameter values where chosen as in Table S1.

The parameters of the CPM model and those of the coarse grained MD simulation are unscaled and require a scaling to fit to measurable data. We follow previous work (Tsingos et al., 2023) for this scaling and we briefly summarize the main points. A single lattice site of the CPM is set equal to 0.25*μ*m *×* 0.25*μ*m, and 10^4^ model timesteps is roughly 8 hours.

As in our previous work, we fit the force units of the model to match up on the widely varying tensile modulus of collagen *E* and we approximated the collagen fibers with *E* = 10^6^Pa which yields a spring constant of 3.1 · 10^−2^Nm^−1^ by approximating a collagen fiber as a cylindrical rod of diameter 0.25*μ*m and applying the formula *K* = *EA/L* with *A* = *π*0.25^2^*μ*m^2^, the cross-sectional area and *L* = 1.5625*μ*m the length of a collagen segment. This choice results in a contraction force of 3.1 · 10^−4^Nm^−1^, leading to typical traction forces ranging from 8nN to 15nN on a single FA, with an total range of 10^−8^ to 10^−10^*N*. These values are slightly lower than those reported in (Tsingos et al., 2023), but fall in the correct order of magnitude of cells traction forces (Wakatsuki et al., 2000; Labernadie and Trepat, 2018).

### 2.6 Statistical significance

The error bars shown in Figure 2-4 denote the mean plus and minus 1 standard deviation. For statistical significance we applied a Welch test. Statistical significance: We computed a *P*-value with the Welch’s test and denoted (*) *P <* 0.05, (**)*P <* 0.01. to compute a *P*-value *P* which we reported with the following symbols: (ns) *P* ≥ 0.05, (*) *P <* 0.05, (**) *P <* 0.01, (***) *P <* 0.001, (****) *P <* 0.0001.

**Figure 2:**
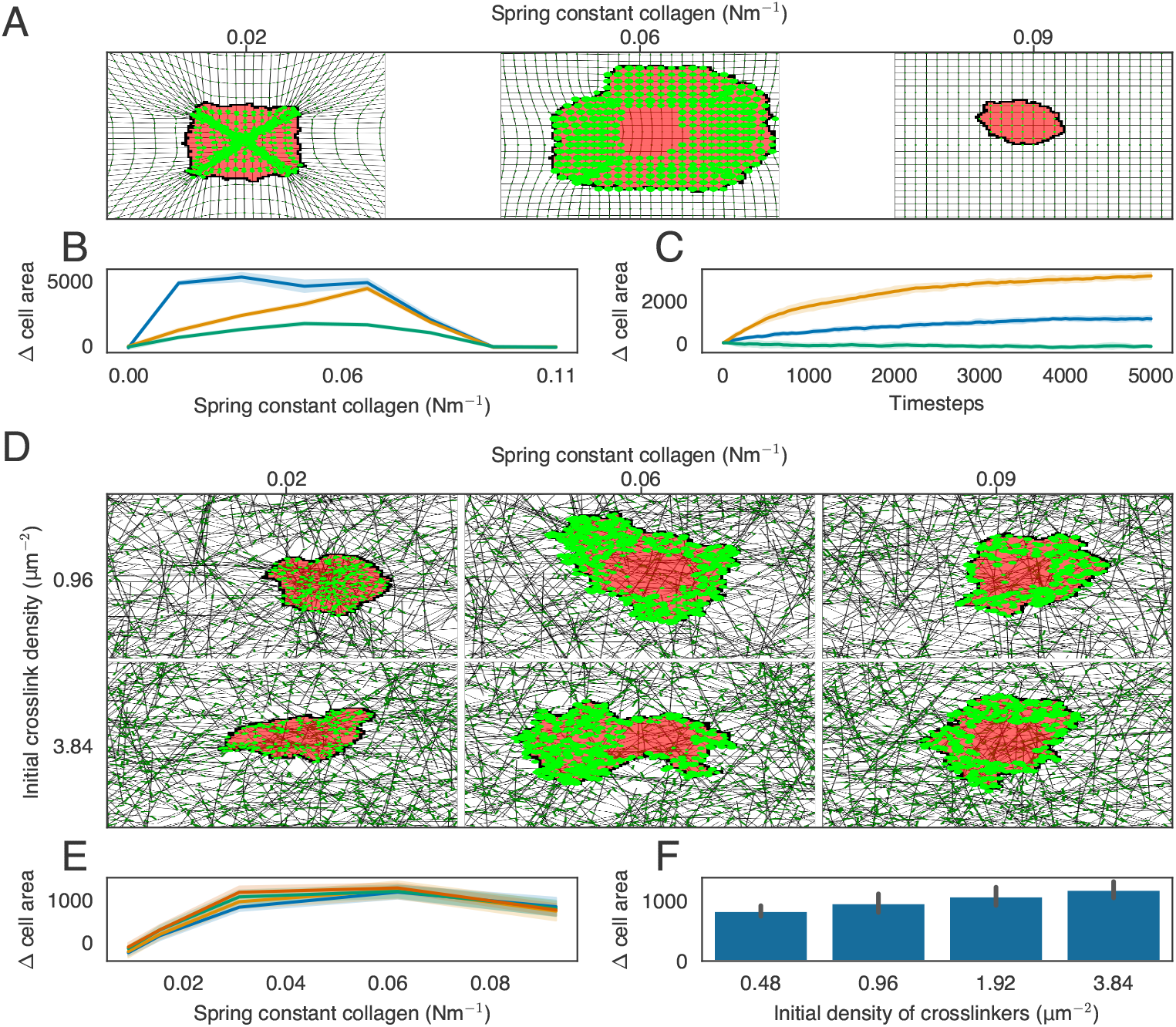
We investigated cell spreading on isotropic ECMs with regular or network structure. A: An example of cell spreading on an isotropic ECM for three different spring stiffness. B: The difference in cell area from the starting size for ECMs of different stiffness. Colors indicate the factor with which we multiply the contraction force of the cell: 1.0 (blue), 5.0 (orange) and 10.0 (green). C: time-evolution of the cell area for different stiffness (blue = 0.015Nm^−1^, orange = 0.047Nm^−1^ and green = 0.09.3Nm^−1^). D: Simulation screenshots of cell spreading on isotropic ECMs with different spring constants and different cross-link densities. E: The difference in cell area from the starting size for ECMs of different stiffness. Colors indicate the cross-link density: 0.48*μ*m^−2^ (blue), 0.96*μ*m^−2^ (orange), 1.9*μ*m^−2^ (green), and 3.8*μ*m^−2^ (red). F: Barplots of final cell size for different collagen densities, using stiffness parameter *K* = 0.031Nm^−1^.

## 3 Results

### 3.1 Stiffness dependent cell spreading

Cell spreading emerges in models of mechanosensitive dynamic FAs with uniform isotropic ECM (Rens and Merks, 2020), suggesting that we should be able to capture cell spreading in the the present model as well. To stay close to the simulated case of the isotropic linear elastic material used previously, we first studied cell spreading on a homogenous matrix constructed by creating long vertical and horizontal strands and cross-linking these at the intersections. On this homogenous matrix, the cells spreading area depends on matrix stiffness in a biphasic manner. Up to an optimum stiffness the cell spreading area increases with matrix stiffness, after which the cell spreading area decreases with matrix stiffness(Figure 2(A,B), Video S1-S2). After this optimum stiffness the tension within the FAs reach the slip regime of the integrins within the FAs. This biphasic effect was not observed in our previous model (Rens and Merks, 2020), or in endothelial cells (Reinhart-King et al., 2005). However, such a biphasic response of cell spreading to matrix stiffness was observed in fibroblasts and T-cells Oakes et al. (2018); Wahl et al. (2019). Interestingly, both the increase in the lifetime of integrin bonds (Oakes et al., 2018), and the increase in integrin ligand density result in a monotonic relationship between substrate stiffness and cell spreading area. To test if our model is consistent with this experimental observation, we doubled the slip parameter *ϕ*_*s*_, which would increased integrin lifetime, and observed that the biphasic effect shifts to higher stiffnesses (Figure S1). We did not study the effect of an increased integrin ligand density on cell spreading, since we have not modelled individual integrins. We next tested how contraction force affects cell spreading. Consistent with experiments showing that inhibition of myosin increases cell spreading (Wakatsuki et al., 2003), in our model we find that cell spreading decreases with an increase in cell contraction force (Figure S2, Video S3).

In Rens and Merks (2020), mechanism for cell spreading, cell elongation and durotaxis based on mechanosensitive FAs where found for a cell on a homogenous, regular ECM. Having implemented a conceptually similar model, we recovered the relationship between ECM stiffness and cell spreading on homogenous, regular ECM. Next, we studied the relation between ECM stiffness and cell spreading on inhomogenous, randomized, but isotropic ECMs. We constructed isotropic randomized matrices as described in Section 2.3 by distributing elastic strands of rougly a cell length and cross-linking them together. We observed a biphasic effect on cell spreading when increasing the spring stiffness of the network (Figure 2(E), Video S4-S5). We also modified ECM stiffness by changing the number of cross-linkers (Figure 2(F)). The cell spreading area also increased with cross-linker density. This effect, however, was only visible for specific values of fiber stiffness (Figure S3), as the tensions in an ECM with soft fibers is still low regardless of cross-linker density and the tension in an ECM with stiff fibers is high regardless of cross-linker density.

### 3.2 Cells elongate on aniostropic matrix due to local anistropy of stiffness cues

In vitro, cell extent along fibers in the substrate (Lai et al., 2012; Friedrichs et al., 2007; Sapudom et al., 2023; Chaubaroux et al., 2015) and we hypothesis that the drive behind this elongation is similar to the mechanism of stiffness dependent cell spreading on a regular, isotropic ECM described in the previous section. We therefore asked how ECM isotropy affects cell spreading. Figure 3(A,B) shows example simulations on inhomogeneous matrices with and without anisotropy. Cells placed on a anisotropic matrices elongate along the axis of aniostropy, provided that the fibers are not excessively stiff (Figure 3(A,C)), and that the network is not overly cross-linked (Figure 3(B,D)). This phenomenon occurs because FAs stabilize rapidly under increased tension. In particular, in ECMs composed of parallel fibers, the difference in tension built up along or orthogonal to the fibers is large as fibers resist extension more effectively than bending. Consequently, FAs stabilize more rapidly in the direction of the bias compared to the orthogonal direction, resulting in observed cell elongation.

**Figure 3:**
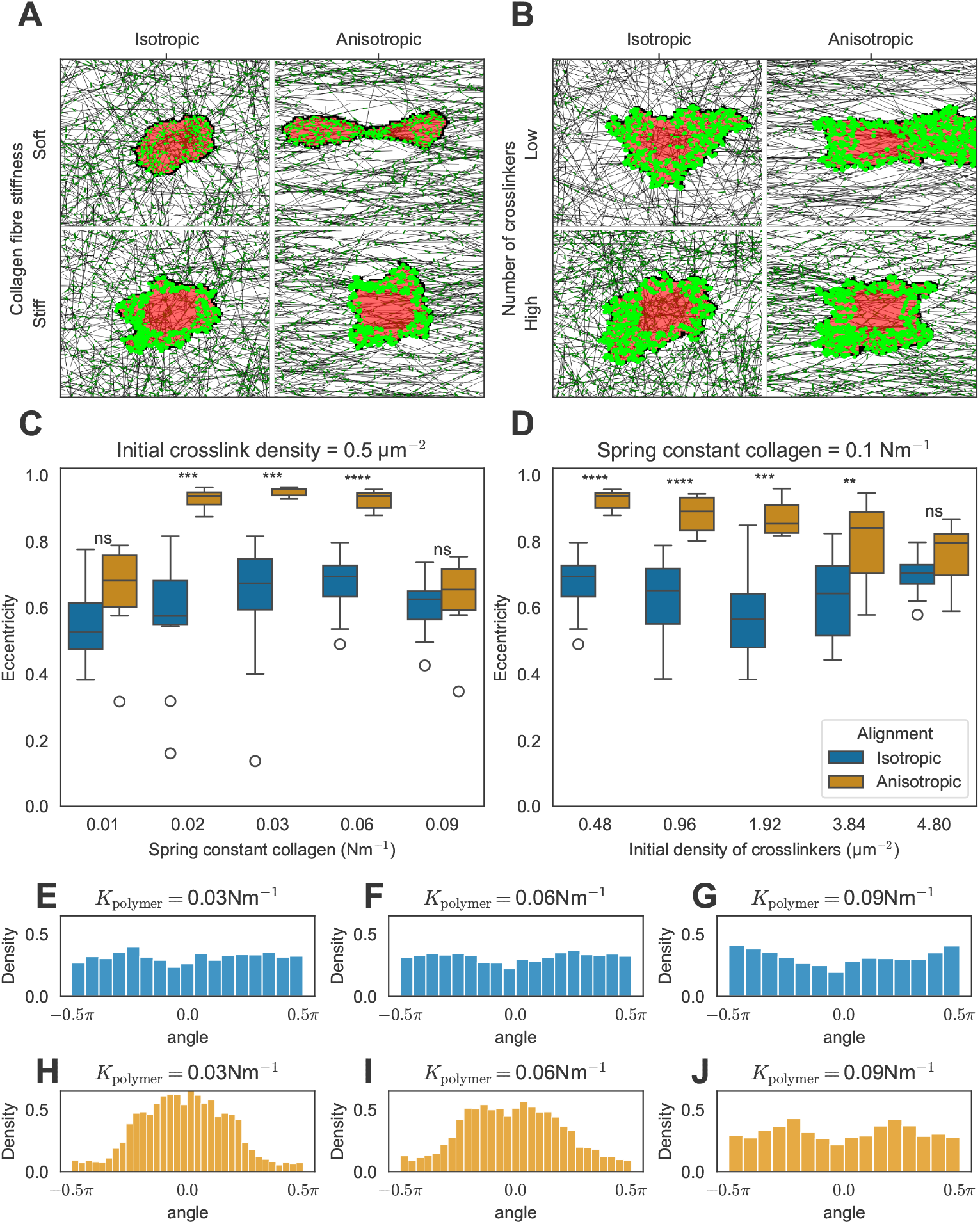
A-B: Screenshots of simulations illustrating the effect of matrix anisotropy on cell shape for different number of cross-linkers and different stiffness of the collagen fibers. C-D: Distributions of the cell’s eccentricity where we increase the ECM stiffness by increasing the collagen spring constant (C) or increasing the number of cross-linkers (D). E-J: Distributions of the angle each FA makes with the horizontal axis that goes through the center of mass of the cell for isotropic ECM (E-G) and anisotropic ECM (H-J).

Consistent with in vitro observations, our model simulations predict that FAs form preferentially at the poles of the cells (Figure 3(A,B)). The angles between the cell elongation axis and the FAs are roughly uniformly distributed on istropic matrices (Figure 3(E-G)). By contrast, on anistropic matrices the FAs are centered on the axis of anisotropy (see single peaks around 0 radians in Figure 3(H,I)). This effect was independent of collagen fibers stiffness, except for the highest stiffness tested where cells no longer elongated and FAs are uniformly distributed around the cell (Figure 3(J)) Similarly, as we increased the degree of cross-linking in anisotropic ECMs with intermediate fiber stiffness the cells failed to elongate (Figure 3(B,D), Video S8-S9). In both cases the fibers provided more resistance to contraction forces perpendicular to the axis of anisotropy, either due to increased fiber stiffness or increased cross-linking. As a result, the tension in the FAs pulling perpendicular to the axis of anisotropy became sufficiently strong for their maturation, leading to rounding of the cell. Interestingly, our model predicts that cells orient perpendicularly to the axis of ECM anisotropy on strongly cross-linked matrices composed of stiff fibers (Figure S4, Video S10), because the FAs at the cell poles enter the slipping regime. Finally, we looked at how the degree of isotropy (as quantified by the order parameter *S*, with *S* = 0 for full isotropy and *S* = 1 for full anisotropy) affects cell shape. The cell eccentricity increased monotonically as a function of the order parameter *S* (Figure S5). In short, we showed that the ECM isotropy influences cell morphology as the ECM geometry determines local perceived stiffness of the cell.

### 3.3 Cell remodels fibers before it starts spreading

In the above we studied how static aniostropies of the ECM can affect cell shape. We next looked at potential cell shape changes due to dynamic, local remodeling of the ECM, potentially driving local ECM anistropies. To measure the local anisotropy of the ECM, we subdivided the domain in square bins of 5 × 5 lattice sites and we compute the order parameter *S*_*i*_(*t*) for the fibers in each bin. Furthermore, we quantified the degree of cell spreading *C*_*i*_(*t*) in a bin *i* as the ratio of the number of lattice sites belonging to the cell and the total number of lattice sites in the bin (i.e., 25). We perform smoothing of the functions *S*_*i*_(*t*) and *C*_*i*_(*t*) by computing a moving time average over the past 10 time steps. Next, we use the functions *S*_*i*_(*t*) and *C*_*i*_(*t*) to study ECM remodeling by the cell.

Figure 4(A) shows two states of a model simulations, one shortly after initialization, and after 10^4^ timesteps showing a large protrusion at the upper right side of the cell. To quantify the degree of cell spreading and ECM alignment in this region, we focused on a square-shaped region around the protrusion of size 20 × 20 lattice sites (i.e., 16 bins) and study the local ECM alignment *S*_*i*_(*t*) and the local cell spreading *C*_*i*_(*t*). An example graph for a single bin is shown alongside the screenshots in Figure 4(A), with the order parameter *S* shown in blue and the degree of cell spreading shown in red. These graphs depict distinct low and high states, with a pronounced transition between them. To examine the variations in the onset of timing of these transitions, we fitted a sigmoid function to each graph:

**Figure 4:**
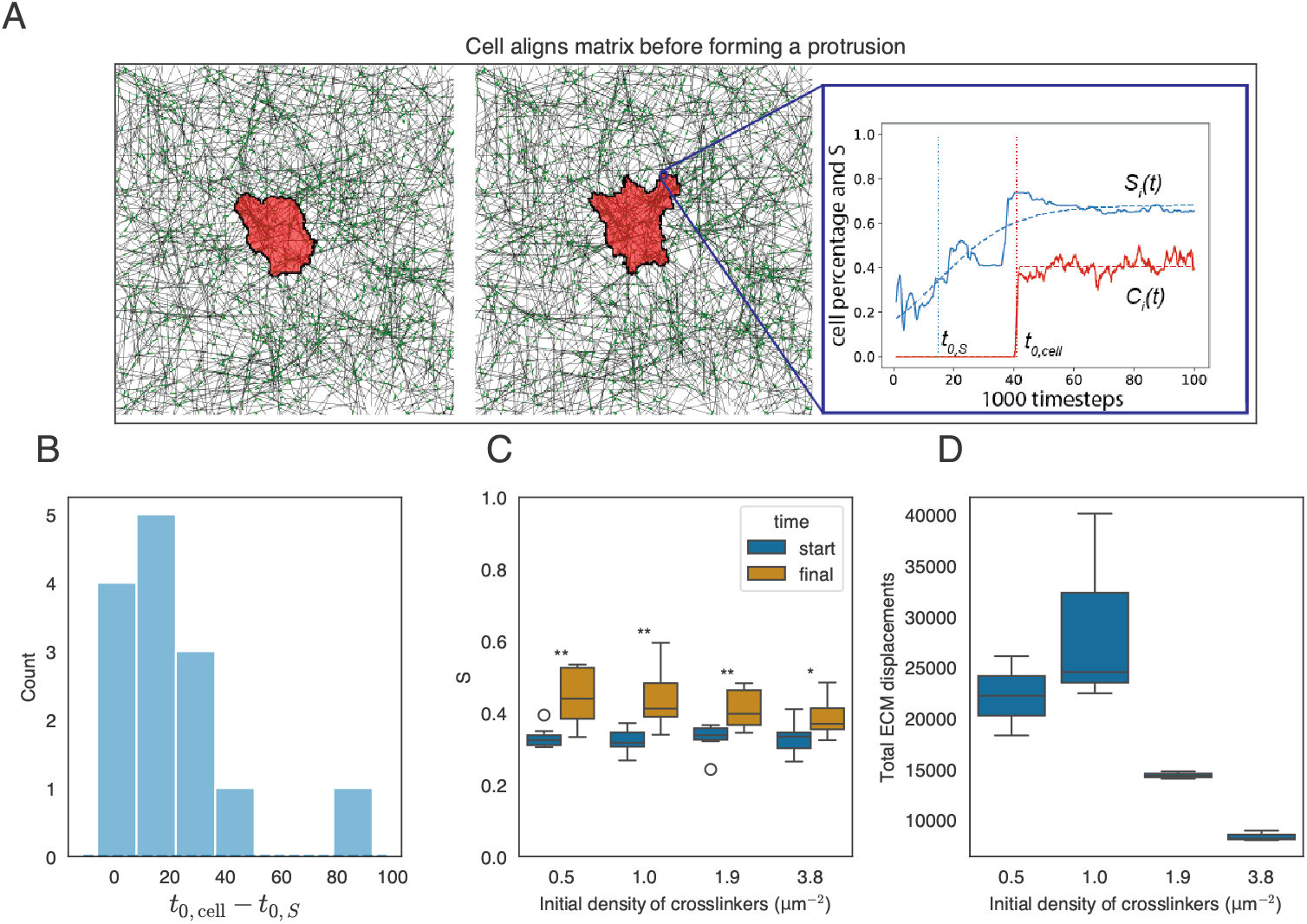
Here we quantified the effects of cell remodeling on the alignment of the ECM and the effect that aligned ECM has back on cell spreading. A: Cell state at beginning of simulation and at the end of simulation. An example of the time series *S*_*i*_(*t*) (red) and *C*_*i*_(*t*) (blue) for a specific bin *i* together with a fitted sigmoid function is included (dashed line) and the center of the fitted sigmoids are indicated on the *x*-axis. B: Distribution of the difference in time of ECM alignment and cell spreading. Almost all mass is on the positive *x*-axis, indicating that the reorientation is happening before the cell spreads. C: Order parameter of a radius around the cell at initial time (pink) and after 10^4^ timesteps (black) for different cross-link densities. D: Total displacements of ECM beads for different cross-linkers densities.

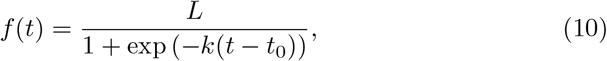

and plotted the distribution of *t*_0,cell_ − *t*_0,*S*_ in Figure 4(B), where *t*_0,cell_ and *t*_0,*S*_ represent the onset times of cell spreading and matrix alignment, respectively. Most of the mass of this distribution is positive, indicating that the cell remodels the matrix first and then spreads over the remodeled fibers.

The cells not only remodeled the ECM at the pseudopodia but also seemed to remodel the ECM all around the cell. We therefore quantified the alignment around the cell by taking the average of order parameters in an annulus around the cell given by

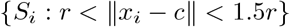

where *S*_*i*_ is the order parameter at the end of the simulation in bin *i, x*_*i*_ is the middle of bin *i, c* is the center of the cell and *r* is the length of the cell. Figure 4(C) shows that the average order parameter around the cell increases over time, and that cross-linking influences the extent of realignment, as higher cross-linker densities decrease the impact of remodeling of the ECM. Additionally, a biphasic effect is seen between the cross-link density and final matrix displacements. Figure 4(D) presents the average displacements of the matrices after 10^4^ timesteps for varying cross-link densities. Fiber networks with a low number of cross-linkers show less displacement compared to those with a medium number. Finally, highly cross-linked fiber networks again show reduced displacements. This biphasic effect on matrix displacement becomes evident when considering two extreme cases: an ECM with few cross-links and an ECM with many cross-links. In the case of an ECM with few cross-links, the displacements induced by the cell do not propagate effectively through the matrix, resulting in low overall displacement. Conversely, in a highly cross-linked ECM, the force required to displace the matrix exceeds the cell’s contractile force, thereby leading to minimal displacement.

## 4 Discussion

We developed a model that dynamically describes cell shape in response to fibrous ECM. We describe cell spreading and cell elongation on fibrous ECM with and without a bias in the orientation. Furthermore, we showed how the cell remodels its surrounding ECM and spreads over parts of this remodeled ECM: The cell pulls on the matrix, reorienting fibers towards itself and then continuously spreading over the newly reoriented fibers.

In our model the relationship between maximum spread area and substrate stiffness is biphasic; the cell spreads more on stifer substrates up to a maximum stiffness after which the maximum area decreases again. This contrasts observations of spreading endothelial cells which monotonically respond to substrate stiffness (Reinhart-King et al., 2005). However, some studies show a biphasic effect for cells when spreading on substrates of varying stiffness (Oakes et al., 2018; Wahl et al., 2019). In the context of this model, the biphasic effect can be explained by focusing on the catch-slip behavior of the integrin types at play, a mechanism that is described before (Oakes et al., 2018). In our model, a FA consists of a cluster of integrins and the unbinding rate of an integrin is the function *d*(*φ*), with *φ* the tension on the integrin. This function determines at which tensions the integrin is detached from the cluster, leading to a smaller FA that is easily detached by the cell. By increasing the parameter *ϕ*_*s*_, that describes the slip-regime of an integrin, integrins will resits larger tensions and hence the cell is also able to spread on stifer substrates. Another idea suggests that not catch-slip bonds are the source of the biphasic effect (as they are in this model), but rather an interplay between different receptors and actin (Wahl et al., 2019). This is an effect that we cannot study with the current model, as the FAs model only take into account a single type of integrin. An more sophisticated contractility and FA models should be implemented to study this.

Next to cell spreading on hydrogels, cell spreading also has been studied on fibrous matrices. Cell spreading was studied on synthetically produced fibrous networks and it was found that increasing fiber stiffness decreases cell spreading (Baker et al., 2015). The authors suggest that this might be caused by higher cell-mediated fiber recruitment for softer fibers leading to more spreading. In a later paper of the same group it was shown that, in a mathematical model, focal adhesions could increase in size on soft fibrous ECMs exactly because of the higher recruitment of possible integrin ligand binding sites (Cao et al., 2017). In our model, an increase in cell mediated fiber recruitment was observed for less cross-linked, hence softer, fibrous ECMs but the positive relation between cell spreading and decreasing stiffness was not found. This discrepancy could be resolved by extending the model in two ways. First, stress dependent crosslinking breakages could be added as it would enhance fiber recruitment for softer ECMs since stress in cross-links is high for sparsely cross-linked ECMs. Secondly, a different FA maturation mechanism that takes into account the availability of nearby fibers should be implemented; currently, higher fiber recruitment increases the number of FAs but does not necessarily increase the size of Fas since the size of a single FA is computed independently of all other FAs.

Our models successfully capture the experimentally observed behavior that cells align to oriented fibers in cross-linked anisotropic networks. Our model suggests that this process follows from the maturation of FAs that sense the stiffer direction of the network. This shows that stiffness-dependent FA maturation suffices for the alignment of cells to oriented ECM. Others have suggested that its not FA maturation which drives this elongation, but rather a positive feedback loop between cell contractility and stresses on the ECM, claiming that the cell increases its contractile force when sensing higher stress (Alisafaei et al., 2022). Contrasting this assumption is the finding that cells do not increase their contractile forces purely based on the stiffness of their environment (Feld et al., 2020). In our explanation of cell elongation, cellular contraction force does not increase on stiffer substrates consistent with the aforementioned observation.

Recently, it has been proposed that the reorienting of fibers by the cell is a two-way process; cell protrusions interact with collagen fibers, to develop anisotropic tension which stabilizes the protrusions in a self-reinforcing cycle (Alisafaei et al., 2022). The mechanical mechanism of tension-dependent FA maturation studied in this paper shows an similar mechanism emerging, where we showed that protrusions are stabilized after the cell aligns surrounding fibers. Regardless of the exact mechanism that drives this mechanical reciprocity between the cell and the ECM, similar mechanical reciprocity plays a role in larger multicellular scales. For example in metastasis (Sander, 2014) as the contractile forces of tumors align surrounding fibers (Balcioglu et al., 2016) and as cancer cells migration is enhanced on aligned fibers forming ‘highways’ (Sander, 2014; Doyle et al., 2022). Conceptually, we implemented a model very similar to the model of Rens and Merks (2020) which models the ECM as a homogeneous isotropic matrix and similar as in Rens and Merks (2020) our model predicts stiffness dependent cell spreading. However, there are discrepancies between the two models: spontaneous cell elongation on substrates of intermediate stiffness and monotonous cell spreading emerge in the model with continuous ECM but were not found in the model presented in this paper. The first discrepancies can partly be explained by the additional assumption made by Rens and Merks (2020), namely that substrate stress strengthens the FAs. The second discrepancy is due to that, in our model, the tensions on the FAs extend into the slip regime, whereas the tension remain lower in the continuous ECM model. This might be remedied with a more realistic contraction model that yields a limited maximum tension on the FAs. There is a final discrepancy between the two models: emergent durotaxis was observed in the model of Rens and Merks (2020), whereas this was not studied in this paper.

In this work we confined our self to modeling a single cell. The presented hybrid model, which combines the versatile cellular Potts model with a fibrous extracellular matrix with dynamic focal adhesions, enables the application of the CPM capabilities. This hybrid model can be utilized to investigate various aspects of the interplay between cells and the extracellular matrix in both singlecell and multicellular contexts. Possible extensions include the study of cell migration along fibers by using one of the many active cell migration models implemented for the CPM such as a polarity vector (Burger et al., 2022), or the Act model (Niculescu et al., 2015); the difficulty lies in the detachment of FAs at the rear of the migrating cell, this could be done, for example, by applying a model for asymmetric traction forces that would rupture the rear FAs or by introducing a chemical symmetry breaking component (Yamaguchi and Knaut, 2022). Another example of possible further study is that of multicellular mechanical interaction. Studies of mechanical cell-cell communication via the ECM have been done using the CPM model (Rens and Merks, 2017; van Oers et al., 2014). These studies, however, applied a linear elastic continuous approach for modelling the ECM, which is a major drawback since linear elastic materials have a low range of mechanical signal propagation. The fibrous ECM model used here is capable of transmitting long-range forces (Tsingos et al., 2023) and when linked in our model of dynamic mechanosensitive FAs could be used in modeling cell-cell mechanical communications. Another extension that is suitable for this model is the inclusion of matrix metalloproteinase which are enzymes that break down the matrix. They can be secreted by cells and are crucial for angiogenesis, tissue remodelling and tissue repair. Current ongoing work involves the inclusion of MMP into the model.

## Author Contributions

KAEK: writing: first draft, coding, conceptualising. ET: conceptualising. RMHM: Funding, conceptualising, writing: editing

## Funding

This work was supported by NWO grant NWO/ENW-VICI 865.17.004 to R.M.H.M. (K.K., E.T., and R.M.H.M.) and by Prof. dr. Jan van der Hoevenstichting voor Theoretische Biologie (R.M.H.M.) affiliated to the Leiden University Fund (LUF). ET is currently funded by the Dutch Research Council (NWO) in the NWO Talent Programme with project number VI.Veni.222.323.

## Acknowledgments

This work was performed using the compute resources from the Academic Leiden Interdisciplinary Cluster Environment (ALICE) provided by Leiden University.

## Supplemental Data

Simulation code is available on request.

## Notes

### Competing Interest Statement

The authors have declared no competing interest.

