## Supplemental Material for "How cells align to structured collagen fibrils: A hybrid cellular Potts and molecular dynamics model with dynamic mechanosensitive focal adhesions"

July 10, 2024

### 1 Supplementary Methods

#### 1.1 Eccentricity and cell shape

To study cell elongation we fit an ellipse to the cell shape. We fit the major axis $A$ and minor axis $B$ of an ellipse such that its moment of inertia is equal to the moment of inertia of the cell shape. The details of this procedure can be found in the supplements of (Rens and Merks, 2017) . The eccentricity of the cell is then defined as

$$e = \sqrt{1 - \frac{B^2}{A^2}}. \tag{1}$$

Clearly, $e \approx 1$ if the cell is elongated since in this case $A \gg B$ . Likewise, $e = 0$ if $A = B$ which is the case when the cell is completely circular.

When the ellipse is fitted, we not only compute the length of the major axis and the minor axis, but also the unit vectors $v_A$ and $v_B$ such that the ellipse is parameterized by

$$c_{\text{cell}} + Av_a \sin(2\pi t) + Bv_B \cos(2\pi t) \tag{2}$$

for $t \in [0, 1)$ and with $c_{\text{cell}}$ the center of the cell.

The vector $v_A$ is used to quantify the direction of the cell elongation. To be more precise, given a unit vector $v$ in a direction to which we want to compare the direction of elongation and the amount of alignment between $v$ and $v_A$ is the absolute value of the inner product between $v$ and $v_A$. We take the absolute value since we do not want to distinguish the directions $v$ from $-v$ nor the directions $v_A$ from $-v_A$.

We quantified the difference in cell area, the number of lattice sites equal to the spin of the cell, from the starting area of the cell which is chosen to be 800 lattice sites.

#### 1.2 Order Parameter

Measuring the alignment of directed objects arises repeatedly in mathematical modeling. Here we use the order parameter to quantify the alignment of collagen strands. Specifically, we are interested in quantifying the alignment of a set of unit vectors $v_1, \ldots, v_m$ in the way of an order parameter. The order parameter is a quantity that is 0 when these vectors point to different directions, and is 1 if they fall on the same line.

A commonly used order parameter is obtained by measuring the angles of these unit vectors with the $x$-axis to obtain angles $\theta_1, \ldots, \theta_m \in [0, \pi)$. One has to be careful when measuring these angles as to obtain angles between 0 and $\pi$ radians, always choosing the minimum of the angle between $v_i$ and either $(1, 0)$ or $(-1, 0)$. Finally, the order parameter $S_{\text{angle}}$ is the modulus of the complex number $\frac{1}{m}\sum_{j=1}^{m} \exp(2\theta_j i)$.

We used a different but equivalent method to compute the order parameter which was inspired by the introduction of a paper on the theory of Q-tensors of liquid crystals (Borthagaray et al., 2020). This method does not require the careful angle measurements as it uses the vectors $v_1, \ldots v_m$ directly. We compute the order parameter $S$ as the largest eigenvalue of the Q-matrix

$$Q = \frac{1}{m}\sum_{j=1}^{m} 2v_i v_i^\top - I, \tag{3}$$

with $I$ the $2 \times 2$ identity matrix.

We will now show that these two methods are equivalent by showing that $S = S_{\text{angle}}$. First we measure the angles $\theta_1, \ldots, \theta_m$ as described above. Next, we consider the vectors $w_i = (\cos(\theta_i), \sin(\theta_i))$ for $i = 1, \ldots, m$. Since $w_i = \pm v_i$ and hence $w_i w_i^\top = v_i v_i^\top$ we compute that $Q$ is of the form

$$Q = \frac{1}{m}\begin{pmatrix} \sum_{j=1}^{m} \cos(2\theta_j) & \sum_{j=1}^{m} \sin(2\theta_j) \\ \sum_{j=1}^{m} \sin(2\theta_j) & -\sum_{j=1}^{m} \cos(2\theta_j) \end{pmatrix}, \tag{4}$$

where we used the standard trigonometric identities $2\cos^2(x) - 1 = \cos(2x) = 1 - 2\sin^2(x)$. The eigenvalue $S$ of $Q$ is then easily recognized, since $Q$ is a symmetric traceless $2 \times 2$ matrix, as

$$S = \sqrt{\left(\frac{1}{m}\sum_{j=1}^{m}\cos(2\theta_j)\right)^2 + \left(\frac{1}{m}\sum_{j=1}^{m}\sin(2\theta_j)\right)^2},$$

which is equal to $S_{\text{angle}}$.

Now we use $S$ in two different ways. To compute the global order parameter $S_{\text{global}}$ and to compute the local order parameter $S_{\text{local}}$. The difference between $S_{\text{global}}$ and $S_{\text{local}}$ stems from the sampling of the unit vectors used to compute them.

The global order parameter $S_{\text{global}}$ is computed as follows. Suppose that $x_1, \ldots, x_n \in \mathbf{R}^2$ are the positions of the beads of the network. Then for each bond $(i, j)$ in the network we define $v_{(i,j)} = \frac{x_i - x_j}{\|x_i - x_j\|}$ and we compute the order parameter using all these vectors as input.

The local order parameter $S_{\text{local}}$ is computed by binning the network in bins of size $l \times l$. Then, for each bin we compute the local order parameter with all bonds that fall in that bin as with the global order parameter. Some bonds are quite long and might cross a bin but might not give a contribution to the order parameter as both end points might lie outside of the bin. To remedy this, we refined the network by interpolating each bond $(i, j)$ in the network with segments of length 1 before computing the local order parameter.

#### 1.3 Measurement of time between onsets

We study locally the the rearrangement of collagen fibers and the spreading of the cell. As the CPM is a discrete model, there is no obvious continuous spreading parameter. We used the binning procedure as in the computation of the local order parameter. We binned the network into square bins of $5 \times 5$ lattice sites. For each of these lattice sites we computed the order parameter $S_{\text{local}}(t)$ over time and we quantified the amount of spreading in that bin as

$$C_{\text{local}}(t) = \frac{\text{number of positive spins in this bin}}{25}. \tag{5}$$

This gives curves $S_{\text{local}}(t)$ and $C_{\text{local}}(t)$ as shown in Figure 4(A) in the main text.

Since we observe that these curves, when the cell makes a pseudopodium, both increase from low to high, we want to quantify the difference in onset of this increase. To this end, we fit sigmoidal functions

$$\sigma_{L,k,x_0}(t) = \frac{L}{1 + \exp(-k(x - x_0))}, \tag{6}$$

to the curves $S_{\text{local}}(t)$ and $C_{\text{local}}(t)$ and we obtain parameter sets $(L_{\text{cell}}, k_{\text{cell}}, x_{0,\text{cell}})$ and $(L_{\text{S}}, k_{\text{S}}, x_{0,\text{S}})$. The difference in onset between $S_{\text{local}}(t)$ and $C_{\text{local}}(t)$ is then the number $x_{0,S} - x_{0,\text{cell}}$ .

### 2 Supplementary Tables and Figures

#### 2.1 Figures

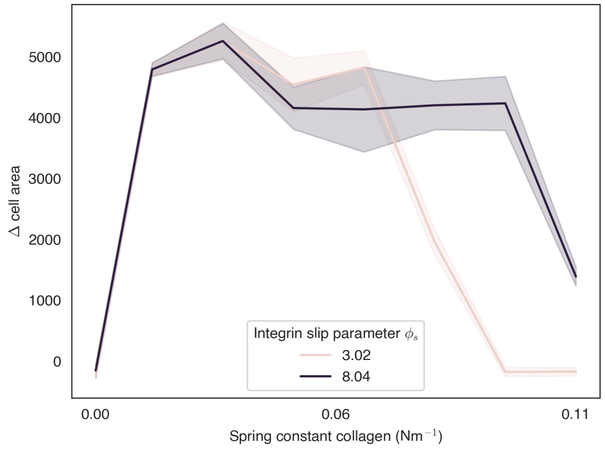


Figure 1: Final cell areas after spreading on a regular ECM with a contraction of 1. The $\phi_s$ parameter describes the slip-regime of the FAs. The cell spreads on stiffer substrates when a higher slip parameter is used.

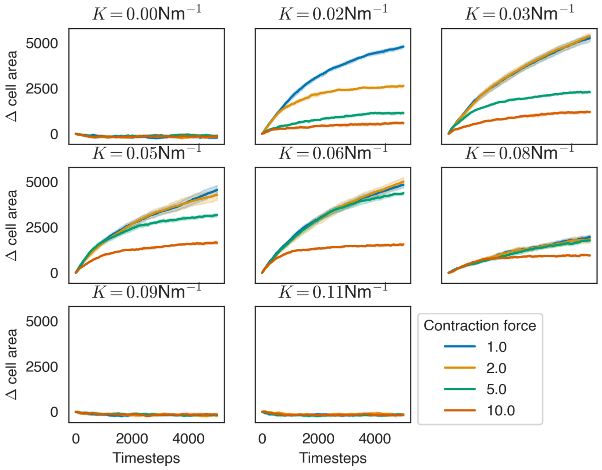


Figure 2: We investigated cell spreading on an isotropic matrix with varying stiffness and with varying cell contraction forces. On soft matrices, we see a large difference between contractile and non-contractile cells with contractile cells spreading the least. This difference decreases when the substrate stiffness is increased and finally disappears when the substrate is too stiff for the cell to spread at all.

| Symbol | Value | Unit | Ref |
|---|---|---|---|
| $\lambda$ | 4.96e+07 | $Nm^{-3}$ | Chosen |
| $J$ | 9.30e-03 | $Nm^{-1}$ | Chosen |
| $A_{\mathrm{ref}}$ | 5.00e+01 | $\mu m^{-2}$ | Chosen |
| $\lambda_{\mathrm{FA}}$ | 800 | – | Chosen |
| $K_{\mathrm{cyto}}$ | 3.10e-04 | $Nm^{-1}$ | Chosen |
| $T$ | 50 | – | Chosen |
| $T_{\mathrm{ECM}}$ | 0.001 | – | (Tsingos et al., 2023) |
| $\gamma$ | 2.88e+00 | $s^{-1}$ | Chosen |
| $d_0$ | 2.88e-02 | $s^{-1}$ | Chosen |
| $f_*$ | 1.29e+10 | $N^{-1}$ | Chosen |
| $\phi_s$ | 4.02 | – | (Novikova and Storm, 2013) |
| $\phi_c$ | 7.76 | – | (Novikova and Storm, 2013) |
| $N_{\mathrm{tot}}$ | 390 | – | (Changede and Sheetz, 2017) |
| $K$ | 3.10e-02 | $Nm^{-1}$ | Chosen |
| $K_{\mathrm{bend}}$ | 3.88e-15 | $N m\mathrm{rad}^{-2}$ | Chosen |
| $\theta_0$ | 3.14e+00 | rad | Chosen |
| Fiber density | 4.80e-01 | $\mu m^{-2}$ | Chosen |
| Fiber anisotropy $\kappa$ | 0, 10 | - | Chosen |

Table 1: Parameter values used in the simulations.

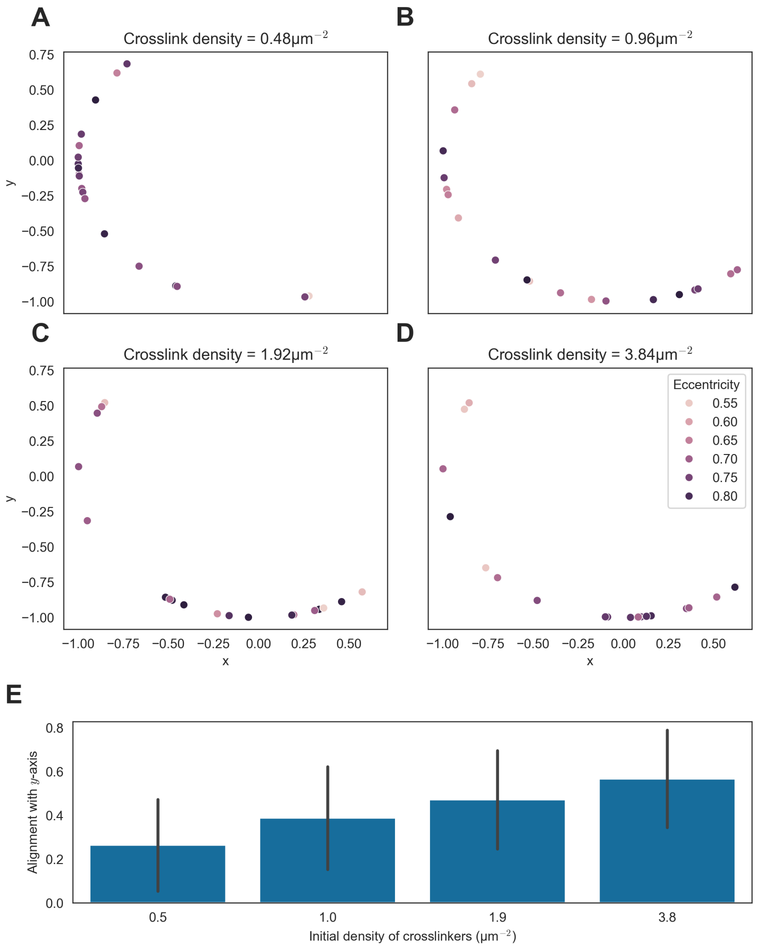


Figure 3: The cell elongates orthogonal to the substrate's direction (the $x$-axis) when the substrate is stiff, here the spring constant is taken to be $0.031\text{Nm}^{-1}$. A-D: The major axis of an ellipse fitted to the cell is shown for different number of cross-linkers. For elongated cells, shown with a darker color, on highly cross-linked matrices, the major axis is close to the y-axis, showing that the cells are aligned to the y-axis. E: A barplot showing the degree of alignment with the $y$-axis for different cross-link densities.

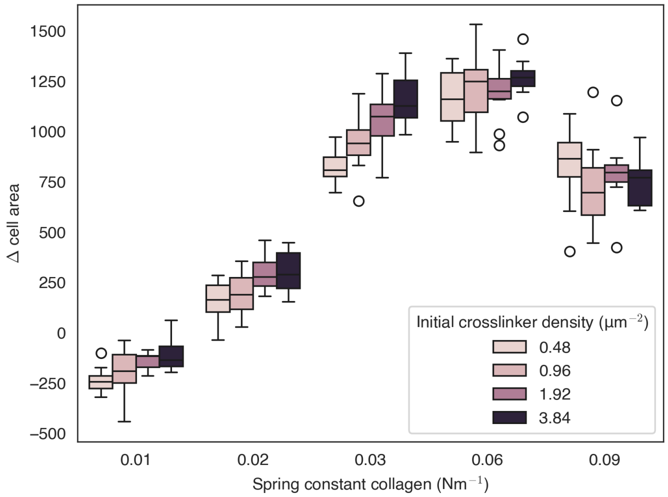


Figure 4: Final cell area distribution for different values of fiber stiffness and different cross-link densities. The effect of cross-linking on the final cell area disappears for very soft and very stiff fibers, but is visible for fibers of intermediate stiffness.

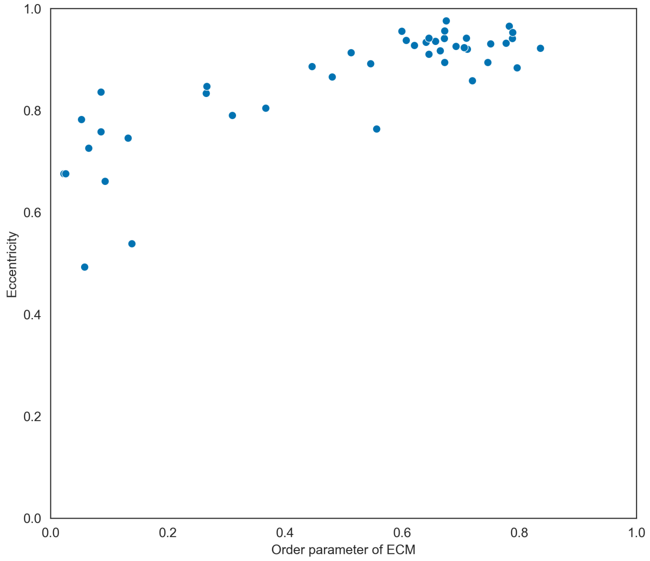


Figure 5: The relation of the global order parameter of the ECM on the final eccentricity of the cell. We see a gradual increasing effect, The relation between the cell's eccentricity and the order parameter of an ECM with a cross-link density of $3.8\mu\text{m}^{-2}$ and fibers of stiffness $0.031\text{Nm}^{-1}$.

**Video S1:** Cell on a regular, soft ECM does not spread fully ($K = 0.016\text{Nm}^{-1}$).

**Video S2:** Cell on a regular, stiff ECM spreads to full size ($K = 0.062\text{Nm}^{-1}$.

**Video S3:** Cell on a regular, stiff ECM spreads to full size ($K = 0.062\text{Nm}^{-1}$, contraction $10\times$ higher).

**Video S4:** Cell spreading on a network-like ECM with soft fibers ($K = 0.031\text{Nm}^{-1}$, crosslink density of $0.96\mu\text{m}^{-2}$).

**Video S5:** Cell spreading on a network-like ECM with stiff fibers ($K = 0.062\text{Nm}^{-1}$), crosslink density of $0.96\mu\text{m}^{-2}$.

**Video S6:** Cell elongating on an anisotropic ECM with soft fibers ($K = 0.031\text{Nm}^{-1}$, crosslink density of $0.96\mu\text{m}^{-2}$).

**Video S7:** Cell does not elongate on an anisotropic ECM with soft fibers ($K = 0.093\text{Nm}^{-1}$, crosslink density of $0.96\mu\text{m}^{-2}$).

**Video S8:** Cell on anisotropic matrix with lower number of cross-linking ($K = 0.062\text{Nm}^{-1}$, crosslink density of $0.48\mu\text{m}^{-2}$).

**Video S9:** Cell on anisotropic matrix with high cross-linking ($K = 0.062\text{Nm}^{-1}$, crosslink density of $4.8\mu\text{m}^{-2}$).

**Video S10:** Cell on anisotropic matrix with high cross-linking and stiff fibers, elongating orthogonal to substrate direction ($K = 0.078\text{Nm}^{-1}$, crosslink density of $4.8\mu\text{m}^{-2}$).
